## Supplemental for "Phenotypic diversity of *Methylobacterium* associated with rice landraces in Northeast India"

**SUPPLEMENTARY TABLES**

**Table S1**: List of distinct *Methylobacterium* isolates sampled in September – October 2016. Sampling states are abbreviated as AR (Arunachal Pradesh) and MN (Manipur).

| SI no. | State | Landrace | Isolate | Closest identified species |
|---|---|---|---|---|
| 1 | AR | Amkil (AMK) | AMKL1 | *Methylobacterium aminovorans* |
| 2 | AR | Amkil (AMK) | AMKS1 | *Methylobacterium radiotolerans* |
| 3 | AR | Amham (AMH) | AMHL1 | *Methylobacterium aquaticum* |
| 4 | AR | Amham (AMH) | AMHS1 | *Methylobacterium radiotolerans* |
| 5 | AR | Ammo (AMM) | AMML1 | *Methylobacterium aerolata* |
| 6 | AR | Ammo (AMM) | AMMS5 | *Methylobacterium aerolata* |
| 7 | AR | Deku (DK) | DKS6 | *Methylobacterium salsuginis* |
| 8 | AR | Deku (DK) | DKL1 | *Methylobacterium p.53* |
| 9 | AR | Deku (DK) | DKL2 | *Methylobacterium rhodinum* |
| 10 | AR | Essum (ESS) | ESSS2 | *Methylobacterium lusitanum* |
| 11 | AR | Essum (ESS) | ESSL2 | *Methylobacterium rhodinum* |
| 12 | AR | Essum (ESS) | ESSS4 | *Methylobacterium salsuginis* |
| 13 | AR | Gegong (GEG) | GEGL2 | *Methylobacterium radiotolerans* |
| 14 | AR | Gegong (GEG) | GEGS2 | *Methylobacterium salsuginis* |
| 15 | AR | Gegong (GEG) | GEGS6 | *Methylobacterium rhodinum* |
| 16 | AR | Gegong (GEG) | GEGS4 | *Methylobacterium extorquens* |
| 17 | AR | Gezang (GEZ) | GEZL2 | *Methylobacterium radiotolerans* |
| 18 | AR | Gezang (GEZ) | GEZS2 | *Methylobacterium radiotolerans* |
| 19 | AR | Dalmang (DAL) | DALS4 | *Methylobacterium radiotolerans* |
| 20 | AR | Dalmang (DAL) | DALL2 | *Methylobacterium salsuginis* |
| 21 | AR | Dalmang (DAL) | DALS6 | *Methylobacterium zatmani* |
| 22 | AR | Nelii black (NELB) | NELBL2 | *Methylobacterium aquaticum* |
| 23 | AR | Nelii black (NELB) | NELBS1 | *Methylobacterium aerolata* |
| 24 | AR | Nelii black (NELB) | NELBS5 | *Methylobacterium komagatae* |
| 25 | AR | Itanagar rice (IT) | ITS5 | *Methylobacterium radiotolerans* |
| 26 | AR | Itanagar rice (IT) | ITS2 | *Methylobacterium salsuginis* |
| 27 | AR | Pallu nelii (PLN) | PLNL1 | *Methylobacterium fujisawaense* |
| 28 | AR | Pallu nelii (PLN) | PLNL2 | *Methylobacterium phyllosphaerae* |
| 29 | AR | Pallu nelii (PLN) | PLNS3 | *Methylobacterium aquaticum* |
| 30 | AR | Pallu nelii (PLN) | PLNS5 | *Methylobacterium aerolata* |
| 31 | AR | Pyapiimepha (PMPHA) | PMPHAS6 | *Methylobacterium fujisawaense* |
| 32 | AR | Pyapiimepha (PMPHA) | PMPHAS1 | *Methylobacterium radiotolerans* |
| 33 | AR | Pyapii (PYA) | PYAL1 | *Methylobacterium komagatae* |
| 34 | AR | Pyapii (PYA) | PYAL2 | *Methylobacterium suomiense* |
| 35 | AR | Pyapii (PYA) | PYAS1 | *Methylobacterium komagatae* |
| 36 | AR | Pangi amm (PAN) | PANS4 | *Methylobacterium radiotolerans* |
| 37 | AR | Pangi amm (PAN) | PANL1 | *Methylobacterium salsuginis* |
| 38 | AR | Rasing (RAS) | RASL2 | *Methylobacterium aerolata* |
| 39 | AR | Rasing (RAS) | RASL1 | *Methylobacterium komagatae* |
| 40 | AR | Taker (TAK) | TAKS2 | *Methylobacterium populi* |

| | | | | |
|---|---|---|---|---|
| 41 | AR | Taker (TAK) | TAKL2 | *Methylobacterium populi* |
| 42 | AR | Tuyim (TUY) | TUYS3 | *Methylobacterium salsuginis* |
| 43 | AR | Tuyim (TUY) | TUYS5 | *Methylobacterium suomiense* |
| 44 | AR | Yagrun (YG) | YGL1 | *Methylobacterium radiotolerans* |
| 45 | AR | Yagrun (YG) | YGS2 | *Methylobacterium radiotolerans* |
| 46 | AR | Yakil (YAK) | YAKL1 | *Methylobacterium salsuginis* |
| 47 | AR | Yakil (YAK) | YAKS1 | *Methylobacterium salsuginis* |
| 48 | AR | Tusing (TUS) | TUSS6 | *Methylobacterium zatmanii* |
| 49 | AR | Tusing (TUS) | TUSS2 | *Methylobacterium komagatae* |
| 50 | AR | Tusing (TUS) | TUSL2 | *Methylobacterium rhodinum* |
| 51 | MN | Abung phou (AP) | APS4 | *Methylobacterium suomiense* |
| 52 | MN | Abung phou (AP) | APS5 | *Methylobaterium aqaumaris* |
| 53 | MN | Abung phou (AP) | APL2 | *Methylobacterium radiotolerans* |
| 54 | MN | Chakhao 1 (CK1) | CK1L1 | *Methylobacterium aquaticum* |
| 55 | MN | Chakhao 1 (CK1) | CK1S3 | *Methylobacterium suomiense* |
| 56 | MN | Chakhao 2 (CK2) | CK2S2 | *Methylobacterium komagatae* |
| 57 | MN | Chakhao 2 (CK2) | CK2L1 | *Methylobacterium suomiense* |
| 58 | MN | Chakhao 3 (CK3) | CK3S3 | *Methylobacterium suomiense* |
| 59 | MN | Chakhao 3 (CK3) | CK3S4 | *Methylobacterium suomiense* |
| 60 | MN | Chakhao pireiton (CKP) | CKPL2 | *Methylobacterium salsuginis* |
| 61 | MN | Chakhao pireiton (CKP) | CKPS1 | *Methylobacterium aquaticum* |
| 62 | MN | Kakcheng phou (KAK) | KAKL1 | *Methylobacterium populi* |
| 63 | MN | Kakcheng phou (KAK) | KAKS3 | *Methylobacterium phyllosphaerae* |
| 64 | MN | Kakcheng phou (KAK) | KAKS1 | *Methylobacterium radiotolerans* |
| 65 | MN | Kumbi phou (KUM) | KUMS6 | *Methylobacterium komagatae* |
| 66 | MN | Kumbi phou (KUM) | KUML2 | *Methylobacterium radiotolerans* |
| 67 | MN | Kumbi phou (KUM) | KUMS5 | *Methylobacterium radiotolerans* |
| 68 | MN | Langphou (LAN) | LANL1 | *Methylobacterium salsuginis* |
| 69 | MN | Langphou (LAN) | LANS1 | *Methylobacterium salsuginis* |
| 70 | MN | Khokngangbi (KHOK) | KHOKL1 | *Methylobacterium suomiense* |
| 71 | MN | Khokngangbi (KHOK) | KHOKS1 | *Methylobacterium komagatae* |
| 72 | MN | Moirangphou (MAN) | MANS3 | *Methylobacterium oryzae* |
| 73 | MN | Moirangphou (MAN) | MANS6 | *Methylobacterium radiotolerans* |
| 74 | MN | Moirangphou (MAN) | MANL2 | *Methylobacterium radiotolerans* |
| 75 | MN | Moirangphou (MAN) | MANL1 | *Methylobacterium salsuginis* |
| 76 | MN | Phouren-mubi (PM) | PML2 | *Methylobacterium radiotolerans* |
| 77 | MN | Phouren-mubi (PM) | PMS2 | *Methylobacterium komagatae* |
| 78 | MN | Phouren-mubi (PM) | PMS1 | *Methylobacterium radiotolerans* |
| 79 | MN | Phou-ngang (PN) | PNS3 | *Methylobacterium persicinum* |
| 80 | MN | Phou-ngang (PN) | PNS6 | *Methylobacterium aquaticum* |
| 81 | MN | Phou-ngang (PN) | PNL1 | *Methylobacterium salsuginis* |
| 82 | MN | Taothabi (TAO) | TAOS1 | *Methylobacterium salsuginis* |
| 83 | MN | Taothabi (TAO) | TAOS5 | *Methylobacterium aquaticum* |
| 84 | MN | Taothabi (TAO) | TAOL1 | *Methylobacterium radiotolerans* |
| 85 | MN | Tolen phou (TP) | TPL1 | *Methylobacterium salsuginis* |

| | | | | |
|---|---|---|---|---|
| 86 | MN | Tolen phou (TP) | TPS5 | *Methylobacterium aquaticum* |
| 87 | MN | Tolen phou (TP) | TPS2 | *Methylobacterium lusitanum* |
| 88 | MN | Akhan phou (AKP) | AKANL1 | *Methylobacterium aquaticum* |
| 89 | MN | Akhan phou (AKP) | AKANS5 | *Methylobacterium salsuginis* |
| 90 | MN | Laiphou (LAP) | LAPL1 | *Methylobacterium salsuginis* |
| 91 | MN | Laiphou (LAP) | LAPS1 | *Methylobacterium salsuginis* |

**Table S2**: List of distinct *Methylobacterium* isolates sampled from seven focal landraces of Manipur in September 2017. All the abbreviations used for landrace is given in brackets after the landrace name.

| Sl.no. | Landrace | Isolates | Closest identified species |
|---|---|---|---|
| 1 | Chakhao (CKP) | CKPL1 | *Methylobacterium salsuginis* |
| 2 | Chakhao (CKP) | CKPL2 | *Methylobacterium aerolatum* |
| 3 | Chakhao (CKP) | CKPL3 | *Methylobacterium aerolatum* |
| 4 | Chakhao (CKP) | CKPS4 | *Methylobacterium salsuginis* |
| 5 | Chakhao (CKP) | CKPL5 | *Methylobacterium gossipicola* |
| 6 | Phouren-mubi (PM) | PML1 | *Methylobacterium suomiense* |
| 7 | Moirangphou (MAN) | MANL1 | *Methylobacterium suomiense* |
| 8 | Moirangphou (MAN) | MANL2 | *Methylobacterium sp.P53* |
| 9 | Moirangphou (MAN) | MANL3 | *Methylobacterium phyllospherae* |
| 10 | Moirangphou (MAN) | MANL4 | *Methylobacterium komagate* |
| 11 | Kumbi-phou (KUM) | KUML1 | *Methylobacterium komagatae* |
| 12 | Kumbi-phou (KUM) | KUMS2 | *Methylobacterium komagatae* |
| 13 | Kumbi-phou (KUM) | KUML3 | *Methylobacterium sp.P53* |
| 14 | Kumbi-phou (KUM) | KUMS4 | *Methylobacterium suomiense* |
| 15 | Kumbi-phou (KUM) | KUMS5 | *Methylobacterium salsuginis* |
| 16 | Kumbi-phou (KUM) | KUMS6 | *Methylobacterium komagatae* |
| 17 | Phou-ngang (PN) | PNS1 | *Methylobacterium indicum* |
| 18 | Langphou (LAN) | LANS1 | *Methylobacterium aerolatum* |
| 19 | Langphou (LAN) | LANL2 | *Methylobacterium komagate* |
| 20 | Langphou (LAN) | LANL3 | *Methylobacterium radiotolerans* |
| 21 | Abung phou (AP) | APL1 | *Methylobacterium sp. 9HR-3* |
| 22 | Abung phou (AP) | APL2 | *Methylobacterium komagatae* |
| 23 | Abung phou (AP) | APS3 | *Methylobacterium salsuginis* |
| 24 | Abung phou (AP) | APL4 | *Methylobacterium sp. P53* |

**Table S3:** List of *Methylobacterium* isolates sampled from landraces, seeds, soil, grasses, and commercial rice cultivated in the Chingarel field, Manipur.

| Sl.no. | Source | Sample | Isolate | Closest identified Species |
|---|---|---|---|---|
| 1 | Soil | Soil | C1 | *Methylobacterium radiotolerans* |
| 2 | Soil | Soil | C3 | *Methylobacterium radiotolerans* |
| 3 | Soil | Soil | C5 | *Methylobacterium radiotolerans* |
| 4 | Landrace | Abung phou | APS4 | *Methylobacterium suomiense* |
| 5 | Landrace | Abung phou | APS5 | *Methylobacterium aquamaris* |
| 6 | Landrace | Abung phou | APL2 | *Methylobacterium radiotolerans* |
| 7 | Landrace | Chakhao Poireiton | CKPL2 | *Methylobacterium salsuginis* |
| 8 | Landrace | Chakhao Poireiton | CKPS1 | *Methylobacterium aquaticum* |
| 9 | Landrace | Phouren mubi | PML2 | *Methylobacterium radiotolerans* |
| 10 | Landrace | Phouren mubi | PMS2 | *Methylobacterium komagatae* |
| 11 | Landrace | Phouren mubi | PMS1 | *Methylobacterium radiotolerans* |
| 12 | Landrace | Moirang-phou | MANS3 | *Methylobacterium oryzae* |
| 13 | Landrace | Moirang-phou | MANS6 | *Methylobacterium radiotolerans* |
| 14 | Landrace | Moirang-phou | MANL2 | *Methylobacterium radiotolerans* |
| 15 | Landrace | Moirang-phou | MANL1 | *Methylobacterium salsuginis* |
| 16 | Seed | Chakhao Poireiton | CK1 | *Methylobacterium radiotolerans* |
| 17 | Seed | Chakhao Poireiton | CK2 | *Methylobacterium radiotolerans* |
| 18 | Seed | Phouren mubi | PM1 | *Methylobacterium radiotolerans* |
| 19 | Seed | Moirang-phou | MP2 | *Methylobacterium radiotolerans* |
| 20 | Seed | Moirang-phou | MP4 | *Methylobacterium radiotolerans* |
| 21 | GR | Grass | GL3.1 | *Methylobacterium rhodinum* |
| 22 | GR | Grass | GL3.2 | *Methylobacterium salsuginis* |
| 23 | GR | Grass | GL3.4 | *Methylobacterium radiotolerans* |
| 24 | GR | Grass | GL2.1 | *Methylobacterium radiotolerans* |
| 25 | HYV-701 | Commercial rice | HYV-2 | *Methylobacterium salsuginis* |
| 26 | HYV-701 | Commercial rice | HYV-4 | *Methylobacterium salsuginis* |
| 27 | HYV-701 | Commercial rice | HYV-1 | *Methylobacterium radiotolerans* |
| 28 | HYV-701 | Commercial rice | HYV-3 | *Methylobacterium salsuginis* |

**Table S4:** List of distinct *Methylobacterium* isolates sampled from the seeds of landraces from Arunachal Pradesh (AR) and Manipur (MN).

| **Sl.no.** | **State** | **Landrace** | **Isolate** | **Closest identified species** |
|---|---|---|---|---|
| 1 | AR | Amham (AMH) | AMH3 | *Methylobacterium salsuginis* |
| 2 | AR | Amkil (AMK) | AMK2 | *Methylobacterium fujisawaense* |
| 3 | AR | Amkil (AMK) | AMK3 | *Methylobacterium fujisawaense* |
| 4 | AR | Gegong (GEG) | GEG1 | *Methylobacterium aminovorans* |
| 5 | AR | Gegong (GEG) | GEG2 | *Methylobacterium aquaticum* |
| 6 | AR | Gezang (GEZ) | GEZ2 | *Methylobacterium radiotolerans* |
| 7 | AR | Yagrun (YG) | YG1 | *Methylobacterium radiotolerans* |
| 8 | AR | Yagrun (YG) | YG2 | *Methylobacterium radiotolerans* |
| 9 | MN | Chakhao poireiton | CKP | *Methylobacterium radiotolerans* |
| 10 | MN | Kumbi-phou (KUM) | KUM1 | *Methylobacterium radiotolerans* |
| 11 | MN | Moirang-phou (MAN) | MP2 | *Methylobacterium radiotolerans* |
| 12 | MN | Moirang-phou (MAN) | MP4 | *Methylobacterium radiotolerans* |
| 13 | MN | Phouren-mubi (PM) | PM1 | *Methylobacterium radiotolerans* |
| 14 | MN | Phou-ngang (PN) | PN | *Methylobacterium radiotolerans* |
| 15 | MN | Phou-ngang (PN) | PN1 | *Methylobacterium radiotolerans* |

**Table S5:** Repeatability of carbon utilization profile of *Methylobacterium* isolates, tested for a random subset of isolates across two experimental blocks.

| Total isolates | Number of mismatches | | | |
|---|---|---|---|---|
| | Glucose | Fructose | Xylose | Sucrose |
| 30 | 1 | 5 | 2 | 0 |

### SUPPLEMENTARY FIGURES

**Fig. S1**: Isolation of *Methylobacterium* sp. in the field using a customized portable enclosure that can be UV-sterilized.

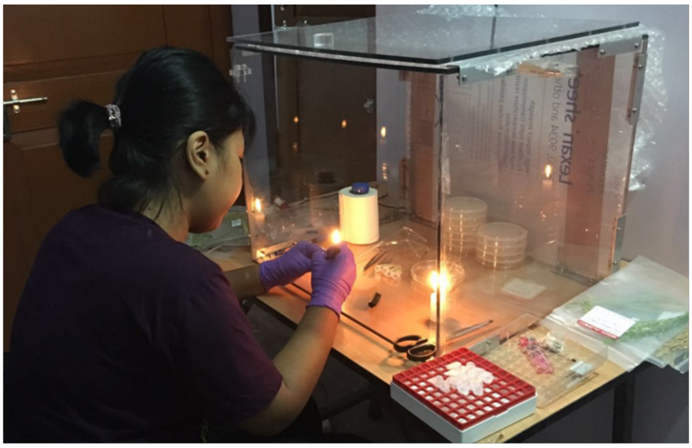

**Fig S2**: **Correlation between geographic distance and phenotypic distance between isolates**. Scatter plot showing pairwise phenotypic distance vs. geographic distance for 91 distinct *Methylobacterium* isolates.

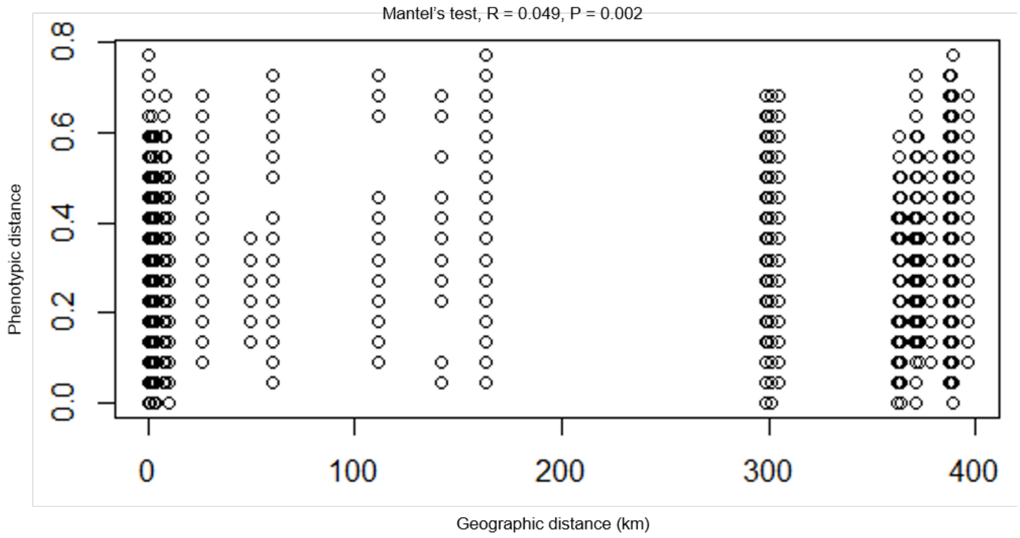

**Fig S3**: **Concentration-dependent growth of *Methylobacterium* isolates on different carbon sources.** (A) A summary of the growth of *Methylobacterium* (58 isolates) on different concentrations of fructose, glucose, xylose, sucrose, and methanol. (B) Fraction of *Methylobacterium* showing different patterns of growth on each carbon source (see key for details). (C) Canonical variance analysis biplots of carbon use profiles, showing the clustering of *Methylobacterium* isolates by sampling state, elevation, and *Methylobacterium* clade.

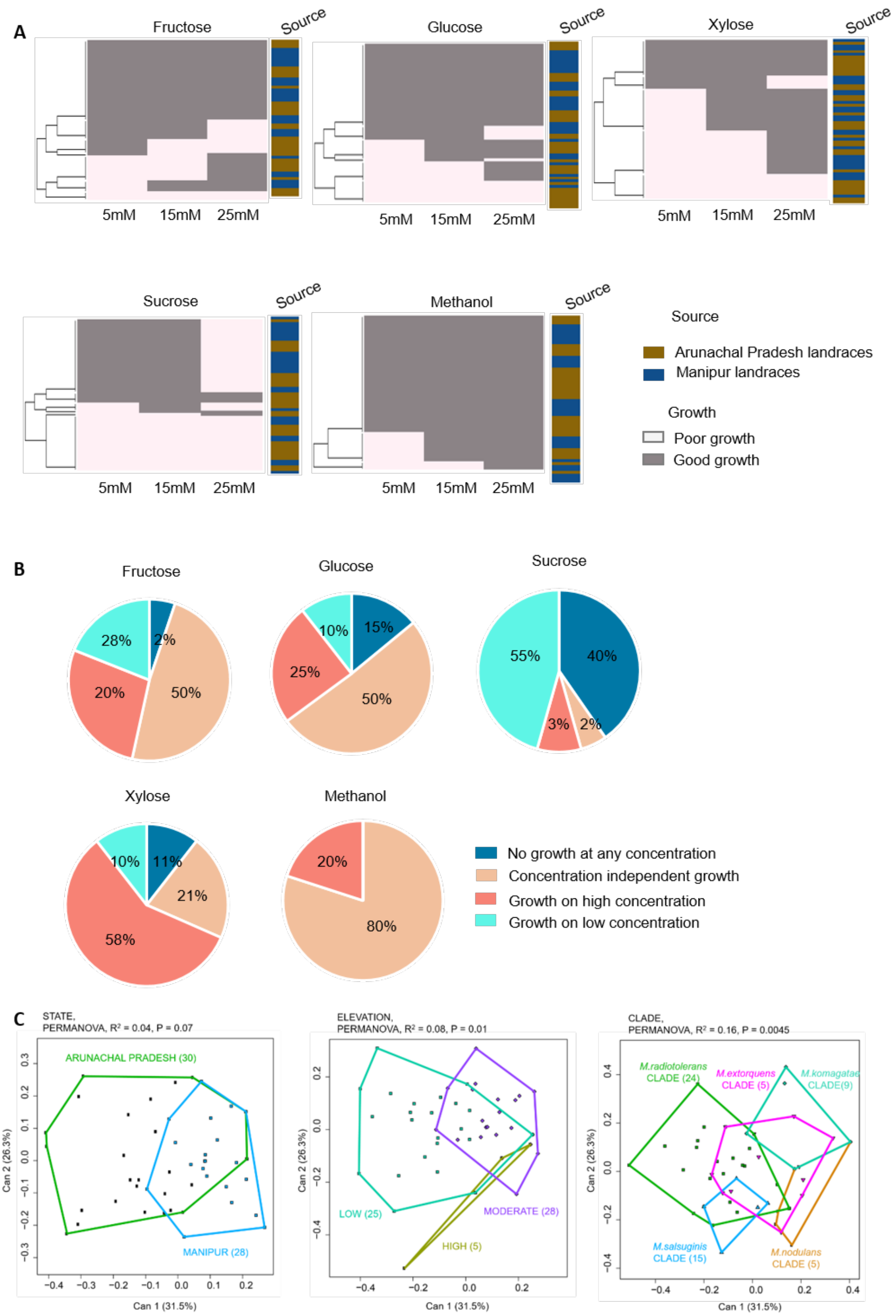

**Fig S4: Density of *Methylobacterium* colonies obtained from various sources.** Samples were imprinted or dilution-plated on Hypho minimal agar plates with 120mM methanol as the sole carbon source. Pink colonies are typical of *Methylobacterium sp.*

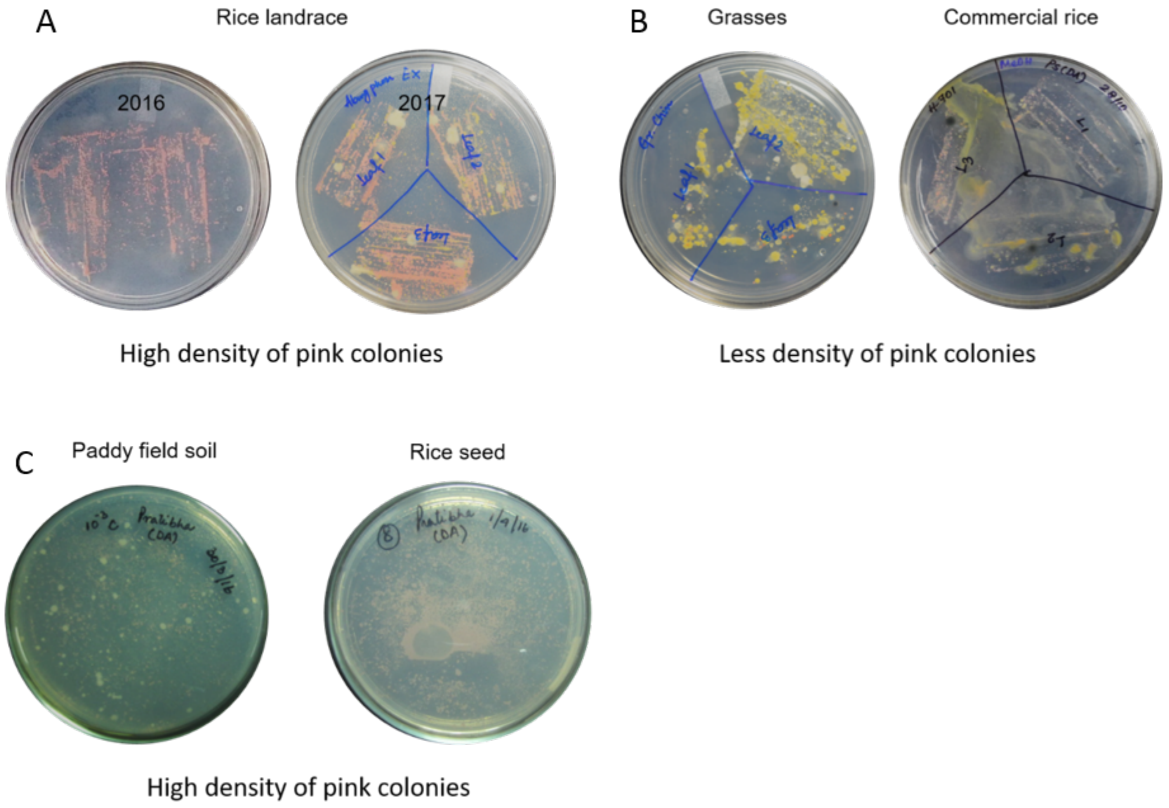

**Fig S5**: ***Methylobacterium* associated with the phyllosphere vs. seed surface of rice landraces**. Each panel shows a pairwise comparison of the *Methylobacterium* community associated with phyllosphere vs. seed for a given each landraces, both in terms of community composition (piecharts) and carbon use phenotype (heatmap). (A) Landraces sampled in Arunachal Pradesh; (B) Landraces sampled in Manipur.

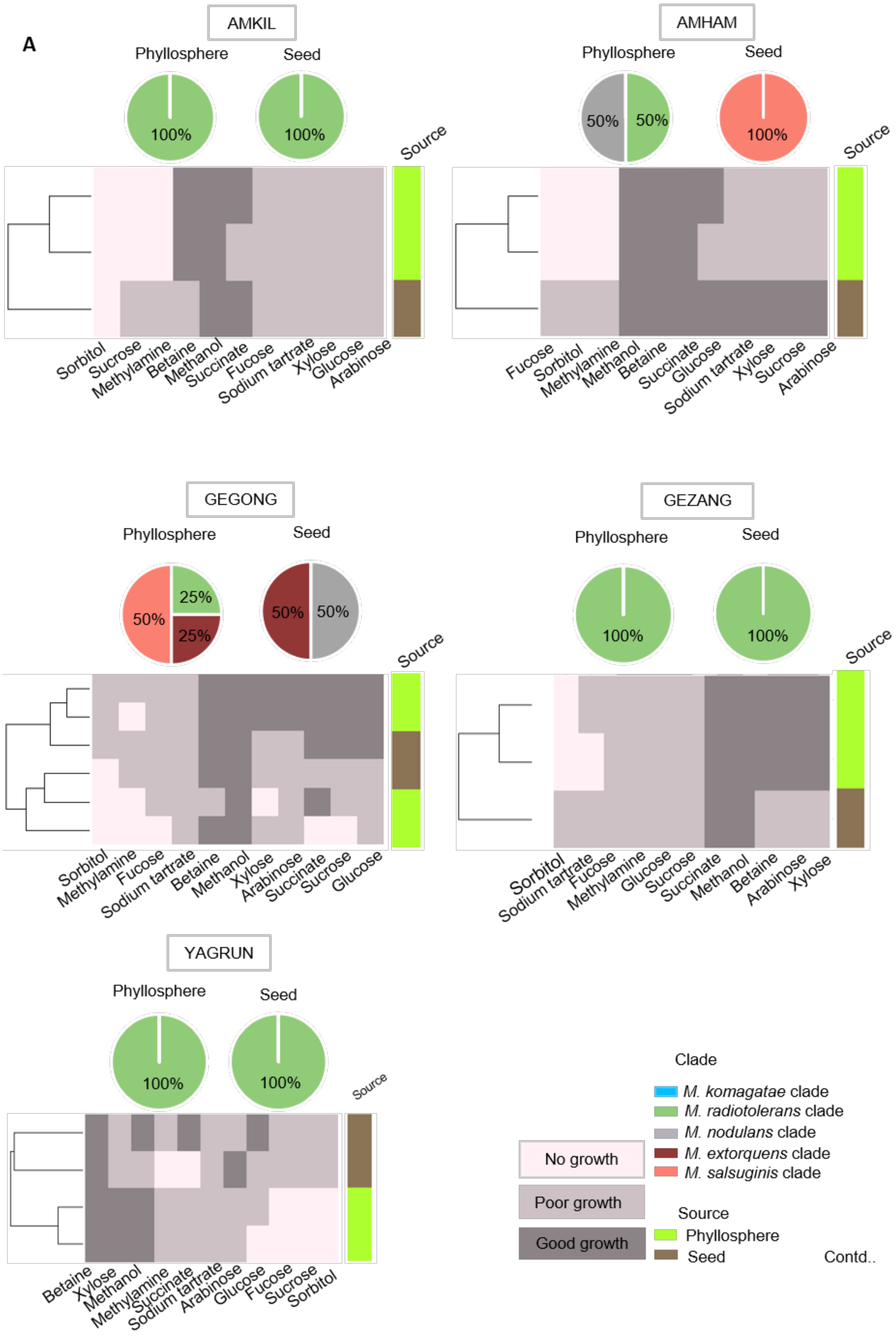


Contd..

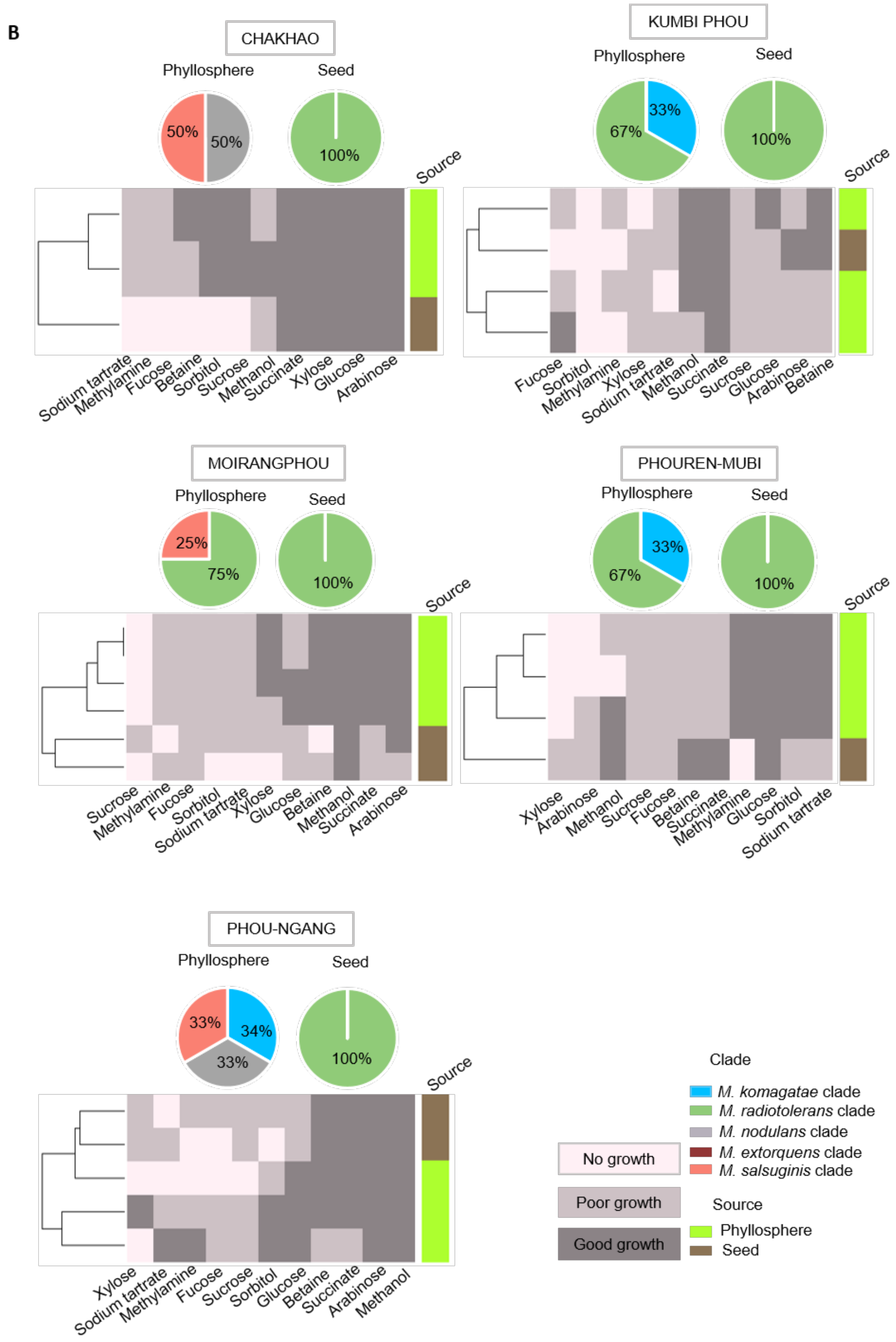
B
CHAKHAO
Phyllosphere
Seed
50%
50%
100%
Source
Sodium tartrate
Methylamine
Fucose
Betaine
Sorbitol
Sucrose
Methanol
Succinate
Xylose
Glucose
Arabinose
KUMBI PHOU
Phyllosphere
Seed
33%
67%
100%
Source
Fucose
Sorbitol
Methylamine
Xylose
Sodium tartrate
Methanol
Succinate
Sucrose
Glucose
Arabinose
Betaine
MOIRANGPHOU
Phyllosphere
Seed
25%
75%
100%
Source
Sucrose
Methylamine
Fucose
Sorbitol
Sodium tartrate
Xylose
Glucose
Betaine
Methanol
Succinate
Arabinose
PHOUREN-MUBI
Phyllosphere
Seed
33%
67%
100%
Source
Xylose
Arabinose
Methanol
Sucrose
Fucose
Betaine
Succinate
Methylamine
Glucose
Sorbitol
Sodium tartrate
PHOU-NGANG
Phyllosphere
Seed
33%
34%
33%
100%
Source
Xylose
Sodium tartrate
Methylamine
Fucose
Sucrose
Sorbitol
Glucose
Betaine
Succinate
Arabinose
Methanol
Clade
M. komagatae clade
M. radiotolerans clade
M. nodulans clade
M. extorquens clade
M. salsuginis clade
No growth
Poor growth
Good growth
Source
Phyllosphere
Seed

**Fig S6: Phylogenetic relationships between rice-associated *Methylobacterium* isolates across years.** Neighbor-joining tree constructed from the 16S rRNA gene sequences of distinct *Methylobacterium* isolates from rice landraces (colored by sampling year), and closely related reference strains (in bold). Bootstrap values ≥ 50% are indicated. Clades are indicated on the right (see Fig 2).

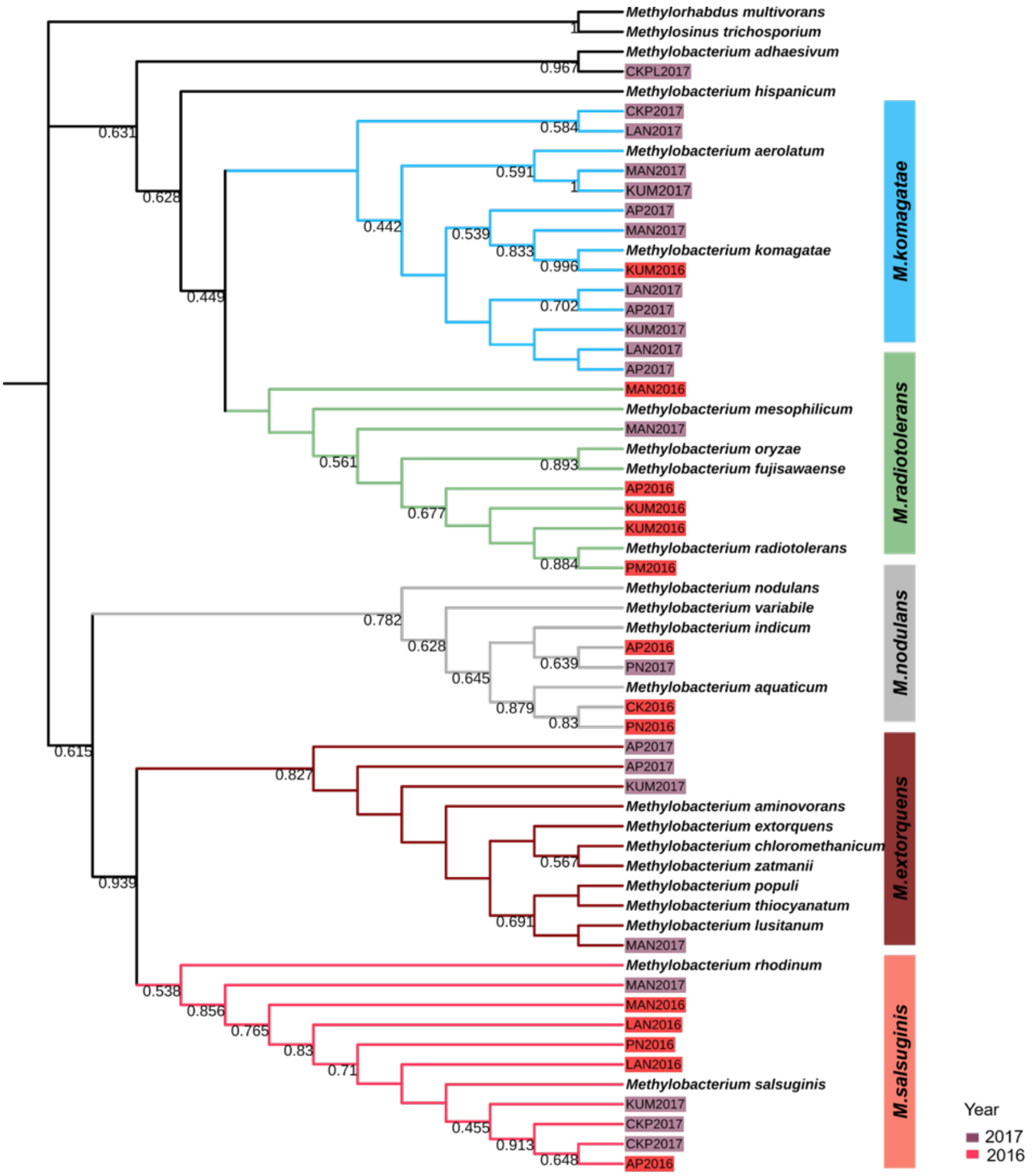

**Fig S7**: **Temporal variation in the composition and carbon use profiles of rice-associated *Methylobacterium* isolates.** Each panel shows a pairwise comparison between bacterial isolates from a given landrace sampled in 2016 vs. 2017 from Manipur, both in terms of community composition (piecharts) and carbon use profile (heatmap).

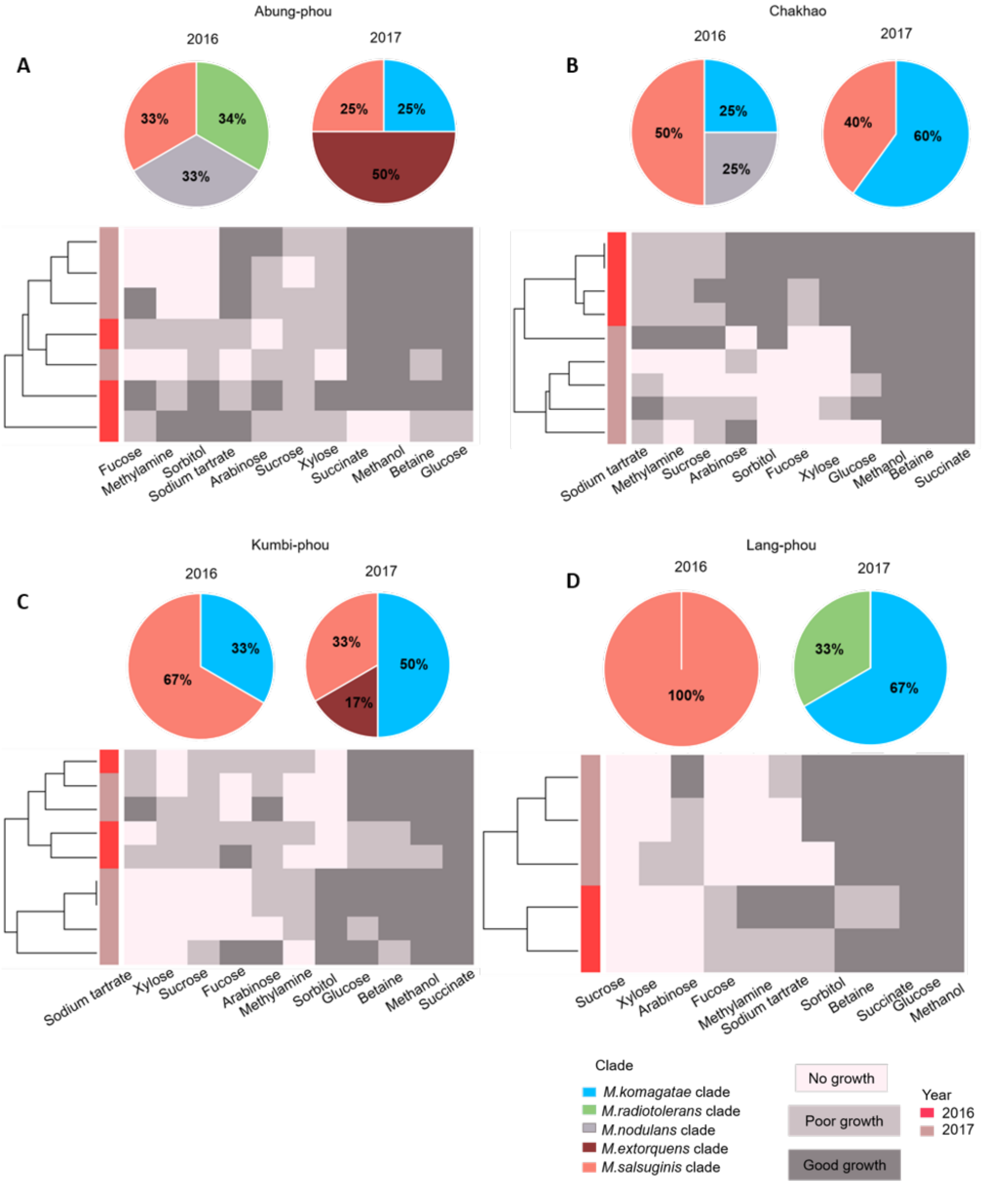


Contd…

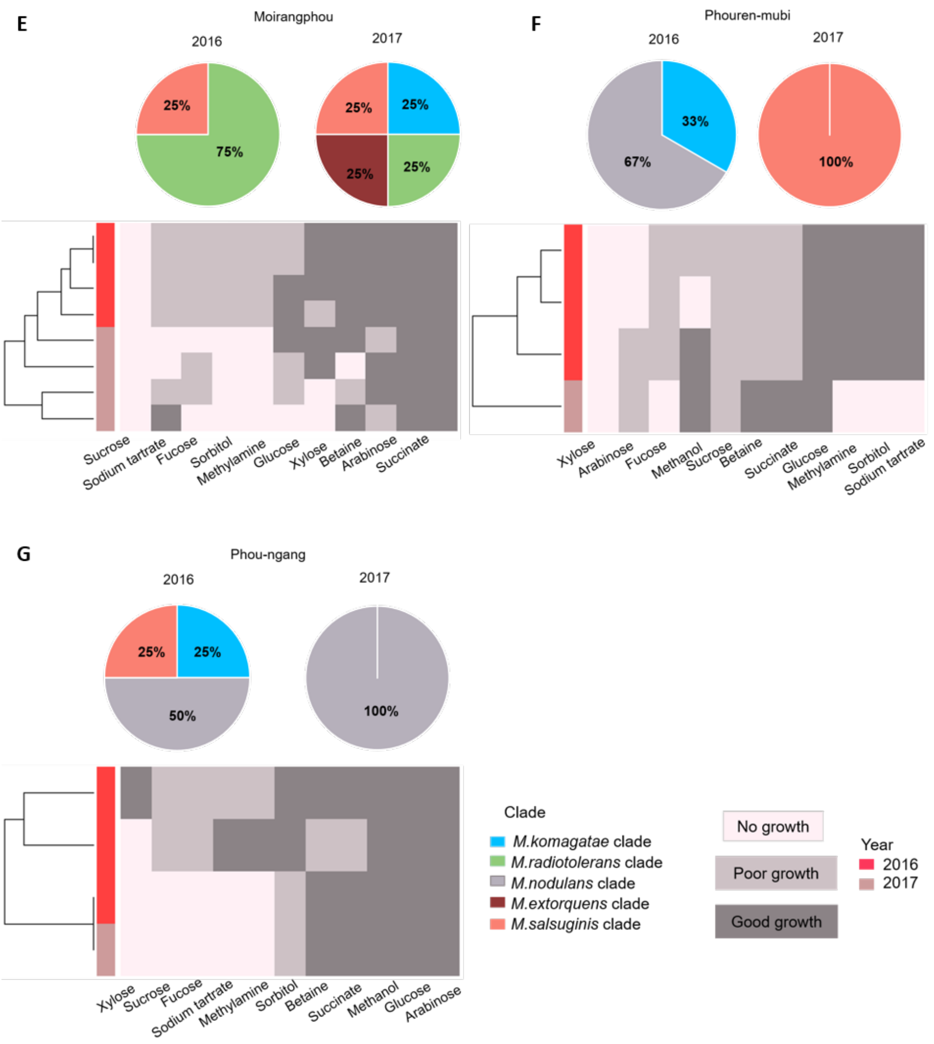

E
Moirangphou
2016
2017
25%
75%
25%
25%
25%
25%
Sucrose
Sodium tartrate
Fucose
Sorbitol
Methylamine
Glucose
Xylose
Betaine
Arabinose
Succinate
F
Phouren-mubi
2016
2017
33%
67%
100%
Xylose
Arabinose
Fucose
Methanol
Sucrose
Betaine
Succinate
Glucose
Methylamine
Sorbitol
Sodium tartrate
G
Phou-ngang
2016
2017
25%
25%
50%
100%
Xylose
Sucrose
Fucose
Sodium tartrate
Methylamine
Sorbitol
Betaine
Succinate
Methanol
Glucose
Arabinose
Clade
M.komagatae clade
M.radiotolerans clade
M.nodulans clade
M.extorquens clade
M.salsuginis clade
No growth
Poor growth
Good growth
Year
2016
2017

**Fig S8: Sugars detected on rice leaf surfaces.** (A) GCMS chromatograms of standard sugars. (B) Representative chromatograms of flag leaf exudates of two landraces: Gegong (Arunachal Pradesh) and Chakhao (Manipur).

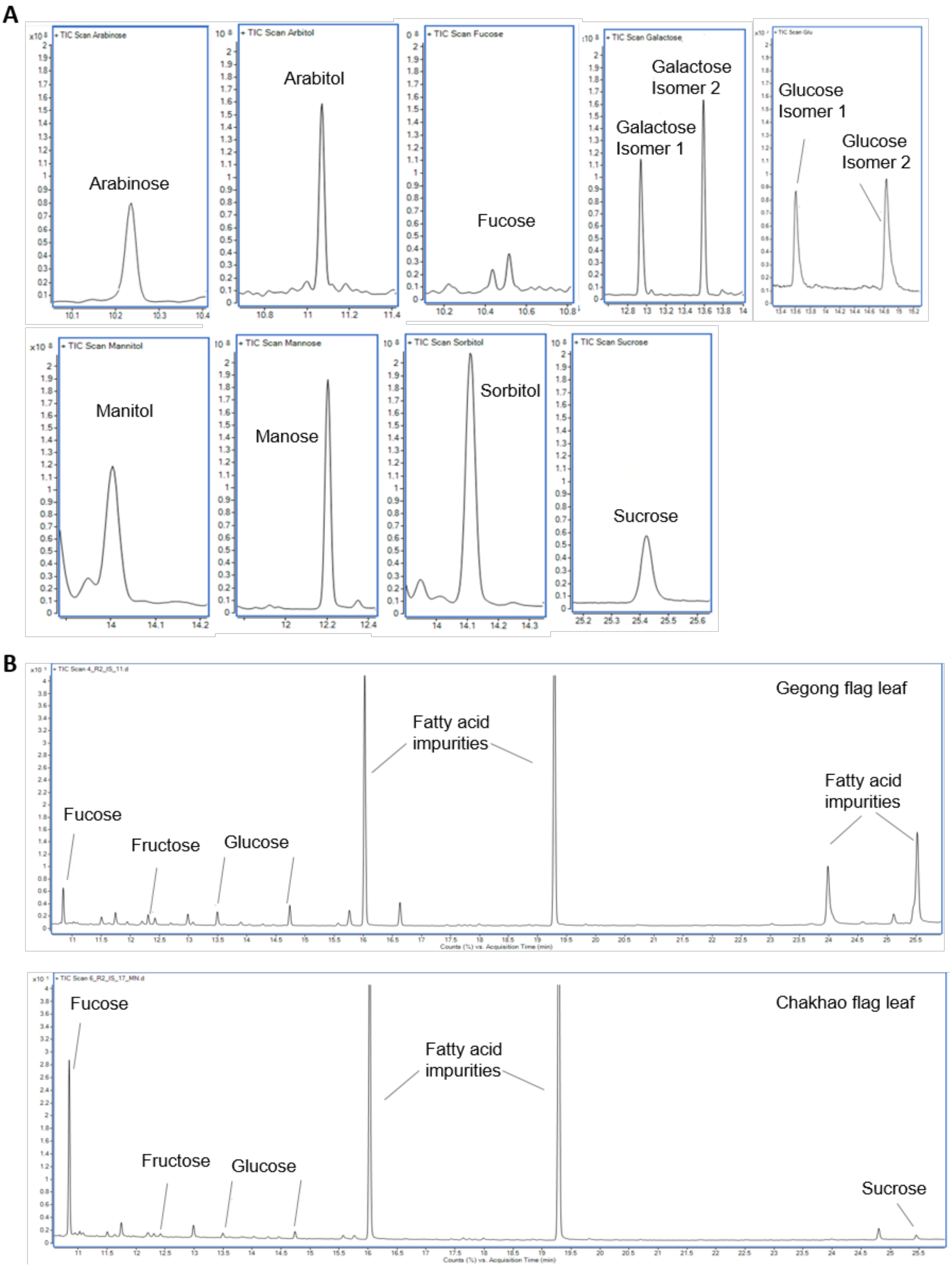
